# Engineering xylose metabolism for diverse polyhydroxyalkanoates synthesis in *Halomonas* TD

**DOI:** 10.64898/2026.05.27.728164

**Authors:** Chang-Le Shen, Bao-Wen Liu, Peng Liu, Yi-Hao Deng, Ling-Ling Zhao, Liu-Song Yu, Wei Situ, Jin-Yan Lv, Hong-Wei Shen, Hai-Tao Yue, Yan-Chun Xiao, Yi-Na Lin, Jian-Wen Ye

## Abstract

Engineering the biosynthesis of fully biodegradable polyhydroxyalkanoates (PHAs) from non-food and renewable feedstocks like lignocellulose is becoming an attractive strategy for sustainable biomanufacturing. However, the efficiency and diversity of PHA synthesis from lignocellulosic hydrolysate (LH), mainly containing glucose and xylose, still remains challenge. Here, *Halomonas* TD, a cost-effective PHA-producing chassis, was developed to utilize glucose and xylose (or LH) for effective production of poly-3-hydroxybutyrate (PHB) and poly(3-hydroxybutyrate-*co*-4-hydroxybutyrate) [P(3HB-*co*-4HB)] by engineering the phosphoketolase pathway-dependent xylose metabolism in the genome, yielding 50.1 g L^−1^ PHB and 42.6 g L^−1^ P(3HB-*co*- 11.4 mol% 4HB) under fed-batch condition. Subsequently, the introduction of Weimberg pathway was found be able to synthesize terpolymer consisting of 3-hydroxybutyrate (3HB), 4-hydroxybutyrate (4HB) and 3-hydroxyvalerate (3HV), namely [P(3HB-*co*-4HB-*co*-3HV)], from glucose and xylose only due to the isoenzyme activity of keto-acid decarboxylase encoded by *kivD*, which converts xylose-derived intermediate 2,5-dioxopentanoate and α-ketoglutarate into butanedial (4HB synthesis) and 2-ketobutyrate (3HV synthesis), respectively. Finally, a tailored-made xylose-induced system was constructed to achieve exquisite xylose transmembrane transportation control for improved synthesis of terpolymer P(3HB-*co*-4.5 mol% 4HB-*co*- 3.0 mol% 3HV), reaching to 6.3 g L^−1^ under shake-flask condition, together with the co-expression of fine-tuned phosphoketolase and Weimberg pathways. This study provides a feasible and sustainable alternative for lignocellulosic resources valorization powered by the engineered *Halomonas* TD capable of efficient xylose utilization and diverse PHAs synthesis.

## 1. Introduction

Polyhydroxyalkanoates (PHAs) is a spectrum of microbial synthesized polyesters, which is known as the most promising environmentally friendly alternative to petroleum-based plastics due to its excellent biodegradability, biocompatibility, and tunable material properties^[1, 2]^. According to the carbon chain length of monomers, PHAs can be classified into three main categories, including short-, medium– and long-chain-length PHA (SCL-/MCL-/LCL- PHA)^[3]^. Currently, the most extensively studied PHAs includes poly-3-hydroxybutyrate (PHB), poly(3-hydroxybutyrate-*co*-4-hydroxybutyrate) (P34HB), poly(3-hydroxybutyrate-*co*-3-hydroxyhexonate) (PHBHHx), poly(3-hydroxybutyrate-*co*-4-hydroxybutyrate-*co*-3-hydroxyvalerate) (P34HBV) and so on. Notably, both P34HB and P34HBV exhibit significant potential in high-end packaging and medical implant application owing to their superior flexibility and thermal stability^[4–10]^. Therefore, in many previous studies, the biosynthesis of P34HB production has been achieved by various metabolically engineered microbial chassis, including *Ralstonia eutropha*^[11, 12]^, *Escherichia coli*^[13]^, etc., using different feedstocks like glucose, soybean oil, fructose, palm sugar, 1,4-butanediol (BDO), γ-butyrolactone (GBL), and so on. Moreover, co-feeding acetate or propionate^[13]^ and engineering cell morphology have been proved to be effective strategies for enhanced P34HB production or increased 4HB molar ratio^[15]^, however, the negative effect on cell growth and PHA accumulation by expressing 4HB synthesis pathway, or supplementing toxic precursor like GBL and BDO, is still an unattainable trade-off for high-yield P34HB production^[16, 17]^. Besides, despite intensive studies on P34HB, the unexpected brittleness resulting from recrystallization (especially for P34HB with low 4HB ratio, < 5 mol%), low transparency and poor thermal stability are still main challenges for widely commercial uses. Recently, the incorporation of 3-hydroxyvalerate (3HV) monomer within P34HB, namely P34HBV, can significantly mitigates the recrystallization behavior with obviously improved transparency and toughness^[18]^ However, to date, the reported production yields of P34HBV in previous studies were pretty lower than some traditional PHAs like PHB^[19–23]^. In addition to the engineering difficulty and production yield limit for P34HB and P34HBV, developing high-performing chassis (e.g., *Halomonas* spp.) able to use renewable non-grain substrate (e.g., lignocellulose-derived sugars) is also an important but challenging strategy to achieve further cost reduction and ecological sustainability^[24]^.

Compared to the fossil fuels, lignocellulosic biomass is an abundant and environmentally sustainable resources that can be converted into value-added chemicals, materials, fuels, etc. through chemical and/or biological processes^[25, 26]^. Particularly, the hydrolysate of lignocellulosic biomass mainly consisting of glucose and xylose is an ideal feedstock for engineered microbes to synthesize various bioproducts including PHAs^[27]^. Generally, most microbial chassis cannot directly use xylose as a carbon source except some microorganisms such as *E. coli*^[28]^, *Saccharomyces cerevisiae*^[29]^, filamentous fungi^[30]^, *Clostridium* spp.^[31]^, *Caulobacter* spp.^[32]^, and archaea^[33]^. Besides, the glucose-induced carbon catabolite repression (CCR) represents a highly prevalent phenomenon across a wide range of chassis cells that generally leads to insufficient utilization of lignocellulosic hydrolysate containing glucose and xylose^[34]^. In these organisms, xylose is typically converted into xylulose-5-phosphate and metabolized via the pentose phosphate pathway (PPP)^[35]^, or cleaved into acetyl-phosphate and glyceraldehyde-3-phosphate by phosphoketolase^[36]^. Recently, metabolic engineering of different microbial chassis has greatly advanced the xylose utilization for the biosynthesis of diverse products including PHB^[37]^, lipids^[38]^, ethanol^[39]^, and BDO^[40, 41]^. Since BDO is a widely used precursor for 4HB monomer accumulation^[17]^, the xylose-to-BDO bioconversion provides a solid foundation for establishing a xylose-to-4HB pathway for 4HB-containing PHAs synthesis (e.g., P34HB and P34HBV).

*Halomonas* species have developed to be superior chassis for low-cost biomanufacturing due to their haloalkaliphilic nature allowing open and non-sterile fermentation process^[42]^. The developments of different genetic parts and tools^[43–45]^ have significantly boosted the engineering capacity and efficiency of *Halomonas* strains to produce various high-value products, including L-threonine^[46]^, ectoine^[47]^, 5-aminolevulinic acid (ALA)^[48]^, and diverse PHAs (e.g., PHB, P34HB^[49]^, P3HBV^[50]^). More importantly, engineering *Halomonas* strains (e.g., *Halomonas* TD) to produce PHB from xylose or lignocellulosic hydrolysates has been proved in previous studies^[51–53]^. Nevertheless, further efforts are still required to expand the diversity of PHA polymers to meet the demands of market sides.

Therefore, this study aims to engineer *Halomonas* TD for efficient biosynthesis of diverse PHAs including PHB, P34HB and P34HBV from glucose and xylose, or lignocellulosic hydrolysate directly.

## 2. Materials and methods

### 2.1 Strains and cultural media

All strains used in this study are listed in Supplementary Table. S1. *Halomonas* TD01, which was isolated from Aydingol Lake of China and stored in CGMCC (China General Microbial Culture Collection Center, Beijing) numbered 4353, was donated by Tisnghua group (Prof. Guo-Qiang Chen). *Escherichia coli* S17-1pir (*E. coli* S17-1) was used as a host for plasmid construction and conjugation into *Halomonas* TD strains.

*E. coli* S17-1 was cultured in Luria-Bertani (LB) medium consisting of 10 g L^−1^ tryptone, 5 g L^−1^ yeast extract and 10 g L^−1^ NaCl. *Halomonas* TD01 and its derivatives were cultured in 60LB medium containing 10 g L^−1^ tryptone, 5 g L^−1^ yeast extract and 60 g L^−1^ NaCl. For agar-plate culture, 1.5% to 2% (wt/vol) of agarose were added in the 60LB medium. The 20LB medium agar plate consisting of 10 g L^−1^ tryptone, 5 g L^−1^ yeast extract, 20 g L^−1^ NaCl and 1.5% to 2% (wt/vol) agarose was used for conjugation by co-culturing the recombinant donor *E. coli* S17-1 and *Halomonas* TD strains. *E. coli* S17-1, *Halomonas* TD01 and their derivatives were all cultured at 37 [. The pH of cultural media was adjusted to 8.0-9.0 using a 5 M NaOH solution for shake flask and fed-batch studies for *Halomonas* TD01 and its derivatives. Furthermore, ampicillin (50 mg L^−1^), chloramphenicol (25 mg L^−1^) and spectinomycin (100 mg L^−1^) were included in the cultural media when necessary to maintain the stability of target plasmids harbored by different recombinant *Halomonas*.

### 2.2 Plasmid construction

The plasmids used in this study are listed in Supplementary Table. S2. Plasmids used in this study were assembled through the Gibson Assembly kit (NEB, Britain) using PCR-amplified DNA fragments, which were purified via Universal DNA Purification Kit (TIANGEN, China). All plasmids were purified using the TIANprep Mini Plasmid Kit (TIANGEN, China). The primer synthesis and DNA sequencing were performed by Songong Co., Ltd. (Shanghai, China). Primer design and sequence alignments were performed by SnapGene software (v6.0.2). Additionally, the genome editing of *Halomonas* TD01 and its derivates was performed using homologous recombination-based method developed in previous studies^[49]^. Genes required to be synthesized or manipulated were listed in Supplementary Table. S3. The ligation products for plasmid construction are firstly transformed into *E. coli* S17-1, thereby colony PCR verification after 12-24 h growth on an agar plate, then the positive clones were sent for sequencing to screen the correct ones for further experiments.

### 2.3 Conjugation

The constructed plasmids were transferred into *Halomonas* strains through a modified conjugation method^[10]^ using *E. coli* S17-1 as a donor cell. The donor cells harboring target plasmids were grown in LB medium with corresponding antibiotics overnight, while the recipient cells were cultured in 60LB medium to reach an OD_600_ of 0.6-0.8. Both of the donor and recipient cells were harvested via centrifugation at 4500 rpm for 2 min (Cence, TGL-16, China), and washed twice with fresh 20LB medium. Subsequently, 50-100 μL of cell mixture (donor-to-recipient volume ratio = 1:1) was dropped onto an antibiotics-free 20LB agar plate for 6-8 h incubation at 37°C. Finally, the conjugated cell lawn was re-suspended using a fresh 60LB medium and spread on a 60LB agar plate containing relevant antibiotics for 48-h incubation to acquire positive colonies with PCR verification.

### 2.4 Shake flask and fed-batch studies

The *Halomonas* TD strains for shake flask fermentation were generally grown on a 60LB agar plate first to obtain single colonies as inoculums for following 12-h inoculation in 5-ml 60LB medium. The pre-cultured seed cells were then inoculated (200 μL, 1 vol%) into 20-ml fresh 60LB medium for 12 h to obtain the 2^nd^ seed cultures. Finally, the 2^nd^ seed culture was 5 vol% (1-mL) added to 20-ml 50MM medium containing corresponding antibiotics, inducers and carbon sources whenever necessary. After 48-h incubation at 37 ℃, 15 mL of cell cultures were harvested via centrifugation at 9000 rpm for 5 min (Cence, H1850, China) for analyzing the cell dry weight (CDW), PHA content and residual concentrations of glucose and xylose. The composition of 50MM includes (g L^−1^): 50 NaCl, 1 yeast extract, 0.5 urea, 0.2 MgSO_4_, 9.65 Na_2_HPO_4_·12H_2_O, 1.5 KH_2_PO_4_, 10 (mL·L^−1^) trace element solution I and 1 (mL·L^−1^) trace element solution II. The composition of trace element solution I is (g L^−1^): 5 Fe(III)-NH_4_-citrate, 2 CaCl_2_ and 1M HCl. The composition of trace element solution II is (mg L^−1^): 100 ZnSO_4_·7H_2_O, 30 MnCl_2_·4H_2_O, 300 H_3_BO_3_, 200 CoCl_2_·6H_2_O, 10 CuSO_4_·5H_2_O, 20 NiCl_2_·6H_2_O, 30 NaMoO_4_·2H_2_O and 1 M HCl.

For fed-batch study, 1-mL overnight seed culture with OD_600_ reaching to 2.5 (± 0.2) was inoculated into 100-mL 60LB medium in a 500-mL shake flask for obtaining 3^rd^ seed culture. Subsequently, 400-mL seed culture was inoculated into a 7-L bioreactor (T&J-Intelli-Ferm B 7L, China) containing 4-L basic cultural medium, which contains (g L^−1^) 50 NaCl, 20 glucose, 10 xylose, 12 yeast extract, 2 urea, 0.2 MgSO_4_, 3.5 KH_2_PO_4_, 40 (mL) trace element I and 4 (mL) trace element II. Three-stage feeding strategy was employed to perform the fed-batch fermentation, including: feeding solution I (Feed-I) containing 267 g glucose and 13 g urea; feeding solution II (Feed-[) containing 333 glucose, 1.3 urea and 2.4 (NH_4_)_2_SO_4_; feeding solution III (Feed-III) containing 400 g glucose only. During the fed-batch fermentation process, Feed-I/II/III were sequentially supplemented into the 7-L bioreactor for maintaining the residual glucose concentration at a range of 5-10 g L^−1^, while the xylose concentration was adjusted from 4 to 8 g L^−1^ by co-feeding a 500 g L^−1^ xylose solution. For fed-batch study using lignocellulosic hydrolysate (LH, Fig. 3E, right panel), which contains 408 g L^−1^ glucose and 174 g L^−1^ xylose purchased from Juwei Yuanchuang Biotechnology Co., Ltd. (Suzhou, China), the Feed-I contained 436 mL LH and 8.7 g urea, Feed-II contained 817 mL hydrolysate, 1.3 g urea and 2.4 g (NH_4_)_2_SO_4_, Feed-III contained LH only. The residual glucose and xylose were off-line measured and monitored hourly using a high-performance liquid chromatography (HPLC, LC-16, Shimadzu, Japan). Cell samples from the fed-batch studies (generally sampled every 4 h throughout the fermentation) were harvested every 4 hours for studying the CDW and PHA content.

### 2.5 CDW and PHA content assays

After 48 h fermentation, cells were harvested via centrifugation at 12000 rpm for 10 min (Cence, H1850, China) and washed once with 20 mL distilled water to generate cell pellet. CDW was determined after 24 h lyophilization of the cell pellet in vacuum freeze-drying treatment. The PHA content of each sample was evaluated by gas chromatography^[54]^ (GC). To determine PHA content, 20-30 mg lyophilized cell powder was sampled and added into a 15-mL tube containing 2-mL chloroform and 2-mL esterification solution (containing 97% v/v methanol, 3% v/v H_2_SO_4_ and 1 g L^−1^ benzoic acid) for 3.5-to-4 h esterification under 100°C for PHA measurement using a GC equipment (GC-2014 Pro, Shimadzu, Japan). In contrast, 15-25 mg PHB (Sigma-Aldrich), 10-15 mg of γ-butyrolactone (Sigma-Aldrich) and 15-25 mg 3-Hydroxyvaleric acid (98% purity, Macklin Biochemical Co., Ltd) were used as standards of 3HB, 4HB and 3HV monomer, respectively.

### 2.6 Measurement of residual glucose and xylose

The residual concentration of glucose of xylose from each fermentation sample were analyzed by a high-performance liquid chromatography (HPLC-20A, SHIMADZU, Japan) equipped with an HPX-87H column (300 mm × 7.8 mm, BioRad). The cell culture was firstly centrifuged at 12000 rpm for 2 min (Cence, H1850, China). The obtained supernatant was diluted 10 times, and filtered with a 0.22 μm filter membrane for HPLC analysis. The mobile phase containing 5 mM dilute sulfuric acid was prepared after 0.22 μm filtration and 30 min sonication. The details for carrying out HPLC analysis were as follows: flow rate, 0.4 ml min^−1^; RID detector; injection volume, 30 μL; column temperature, 55 [. The standard curves of glucose and xylose were generated using a spectrum of sugar concentration: (g L^−1^) 0.6, 1.2, 1.8, 2.4, 3.

### 2.7 Fluorescent Intensity Characterization

The secondary seed culture of different recombinant *Halomonas* strains harboring sfGFP expression vessels was inoculated (1 vol%) into the fresh 50MM medium supplemented with relevant antibiotics and varying xylose concentrations for 12-h cultivation in a 96-deep-well plate. For *E. coli* recombinants, LB medium was used a culture medium instead of 50MM. After 12 h cultivation, the obtained cell cultures were diluted with PBS buffer to an OD_600_ of 0.2-0.8. Followed by simultaneous measurement of fluorescence intensity (FI) and OD_600_ using a microplate reader (excitation/emission: 488/520 nm, Varioskan Flash, Thermo Scientific),the normalized fluorescence intensity level (FI/OD_600_) could be determined.

## 3. Results and Discussions

### 3.1 Engineering xylose utilization by introducing phosphoketolase pathway

Before the introduction of the phosphoketolase pathway, the pre-test of cell growth and sugar consumption by *Halomonas* TDB strain (Fig. 1A), the starting recombinant host, was firstly carried out to study its native xylose utilization capacity. The recombinant TDB, which was derived from *Halomonas* TDH4^[55]^ harboring chromosome-carried *dhaT*-*aldD* (locus G51) and *orfZ* (locus G4) modules able to synthesize PHB and P34HB (when using 1,4-BDO as a precursor), was cultured in 50MM medium supplemented with different carbon sources including 20 g L^−1^ xylose (20X), 20 g L^−1^ glucose (20G) and the mixed one thereof (20G+20X) for 48 h in a 500-mL shake flask (Fig. 1B). The results revealed that the recombinant TDB cannot grow on xylose only in 20X group with no significant growth of OD_600_ and CDW, and negligible consumption of xylose. By contrast, the ‘20G’ and ‘20G+20X’ groups that contains 20 g L^−1^ glucose were able to accumulate 8.5 g L^−1^ and 7.3 g L^−1^ CDW containing over 60 wt% PHB content (Fig. 1B-1D), however, the supplementation of xylose showed obvious negative effect on cell growth, especially for the stationary phase (Fig. 1B-1C). In conclusion, the TDB strain cannot use xylose as a sole substrate unless further metabolic modification. Therefore, after analyzing the annotated genome^[51]^, an IPTG-induced *xylA-xfp* module driven by P_MmP1_ promoter was constructed on pSEVA321, namely P1 (Fig. 1E), in TDB strain to achieve complete phosphoketolase pathway aiming to convert xylose into acetyl-CoA allowing for cell mass and PHB synthesis. Shake flask study was carried out by recombinant TDB harboring P1 construct (TDB-P1) grown on 20X and 20G+20X condition in the presence of 5, 10 and 200 mg L^−1^ IPTG, respectively. Expectedly, the CDW accumulation and xylose consumption levels of TDB-P1 grown on 20 g L^−1^ xylose (20X) increased significantly along with the increased IPTG dosage compared to the control group using start host TDB (Fig. 1F). Similarly, the CDW, PHB content and xylose consumption of TDB-P1 grown on mixed carbon source (20G+20X) under the induction of 200 mg L^−1^ IPTG exhibited over 40%, 17% and 270% increase, respectively, in contrast to TDB group as a control (CDW: 10.6 g L^−1^ *vs* 7.6 g L^−1^; PHB content: 77.9 wt% *vs* 66.1 wt%; xylose consumption: 3.4 g L^−1^ *vs* 0.9 g L^−1^) (Fig. 1G). Moreover, the P34HB synthesis was also assessed by TDB-P1 grown on glucose and xylose (20G+20X) supplemented with 200 mg L^−1^ IPTG, and 2.5 and 5 g L^−1^ 1,4-BDO, which is used as a precursor for 4HB monomer synthesis (Fig. 1H). Surprisingly, in addition to the obvious improvement of cell growth (CDW), the 4HB molar ratio of P34HB synthesized by TDB-P1 was significantly increased up to 9.6 mol% (2.5 g L^−1^ 1,4-BDO) and 13.1 mol% (5 g L^−1^ 1,4-BDO) compared to only 3 mol% by TDB strain grown on glucose under the same condition except removing the IPTG and xylose from the medium (Fig. 1I). These results demonstrate proven feasibility of engineering *Halomonas* TD to using xylose as a feedstock for different PHA synthesis.

**Fig. 1.**
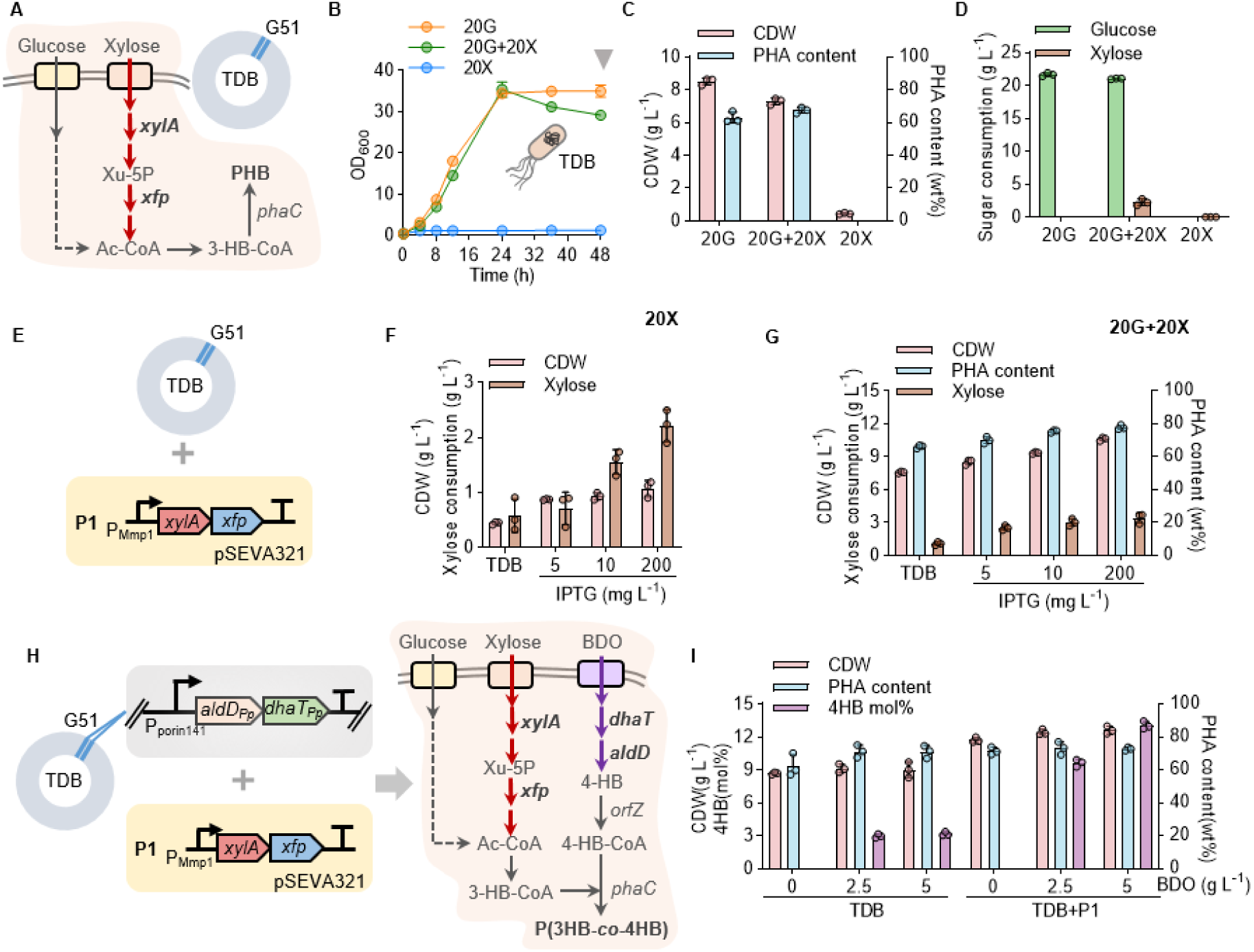
Engineering xylose utilization in recombinant *Halomonas* TDB by introducing phosphoketolase pathway. (**A**) Metabolic pathway design for the co-utilization of glucose and xylose to support the cell growth and PHB synthesis of recombinant TDB, a derivate of *Halomonas* TDH4^[46]^, harboring *dhaT*-*aldD* module controlled by P_porin141_ on chromosome (loci G51). The bioconversion pathway from xylose to acetyl-CoA encoded by *xylA* and *xfp* genes was indicated by bold arrows in red. (**B**) Comparative analysis of the cell growth (OD_600_) by TDB grown on different carbon sources in 50MM medium during the 48-h shake-flask cultivation. Carbon sources include: 20G (20 g L^−1^ glucose, orange line); 20G + 20X (20 g L^−1^ glucose + 20 g L^−1^ xylose, green line); 20X (20 g L^−1^ xylose, blue line). (**C, D**) Cell dry weight (CDW), PHA content, and the consumption of glucose and xylose after 48-h shake-flask fermentation (inverted triangle in gray from part **B**). (**E**) Schema of recombinant TDB containing IPTG-induced *xylA*-*xfp* cluster enabling xylose utilization, namely P1. (**F, G**) CDW, PHA content and xylose consumption by recombinant TDB harboring P1 construct grown on 20 g L^−1^ xylose (20X in **F**) and mixed carbon source (‘20G + 20X’ in **G**), respectively, in the presence of different IPTG concentrations (5, 10, 200 mg L^−1^) in contrast to the start host TDB grown in the same condition (TDB group). (**H**) Schema for P(3HB-*co*-4HB) (P34HB) production by recombinant TDB harboring P1 construct from glucose, xylose and BDO, a precursor for 4-hydroxybutyrate monomer synthesis (4HB-CoA). (**I**) CDW, PHA content and 4HB molar ratio by recombinant TDB with or without P1 construct grown on mixed carbon source (20G + 20X) supplemented with different BDO concentration in the presence of 200 mg L^−1^ IPTG. Detailed descriptions of key enzymes and metabolites were: *xylA* encoding xylose isomerase; *xfp* encoding phosphoketolase; *dhaT* encoding 1,4-butanediol reductase; *aldD* encoding aldehyde dehydrogenase; *orfZ* encoding CoA transferase; *phaC* encoding PHA synthase; Xu-5P, xylulose-5-phosphate. Error bars represent standard deviations, n = 3.

In many microorganisms, the metabolisms of xylose and glucose are incompatible, namely carbon catabolite repression (CCR)^[36]^, which cannot allow sufficient synergetic utilization of glucose and xylose. To assess whether the CCR effect exists *Halomonas* TD when introducing the phosphoketolase pathway, shake-flask study by TDB and its recombinant TDB-P1 grown on 20G+20X and 20G+10X, respectively, was conducted. OD_600_ and residual sugar concentration were recorded at 0, 8, 16, 24, 36 and 48 h during the fermentation process. In contrast to the TDB groups unable to use xylose, TDB-P1 recombinant can synchronously utilize glucose and xylose to achieve higher level cell growth (OD_600_ & CDW) and PHB content without CCR effect (Figs. 2A and 2B, Supplementary Fig. S1). This is probably attributed to the glucose metabolism independent of phosphotransferase system (PTS) in *Halomonas* TD. The consumption of xylose maintained at around 4-to-5 g L^−1^ for both mixed carbon source conditions, while glucose was almost completely consumed. Moreover, the fine-tuned *xylA-xfp* module under the control P_porin140_ promoter, similar transcription level of P_MmP1_ under 200 m g^−1^ IPTG induction (Fig. 1G), was integrated on G53 locus in TDB strain to obtain a chromosomally engineered recombinant (TDBX) allowing robust fermentation test under antibiotic-free condition (Fig. 2C). Fed-batch studies by TDBX were thus carried out in a 7 L bioreactor to achieve PHB (‘- BDO’ group without BDO addition) and P34HB (‘+ BDO’ group with BDO addition) synthesis from glucose and xylose. After 44 h fermentation, over 50 g L^D¹^ PHB (CDW, 74.5 g L^−1^; PHB content, 67.2 wt%) and 45 g L^−1^ P(3HB-*co*- 6 mol% 4HB) (CDW, 66.3 g L^−1^; P34HB content, 68.2 wt%) were obtained (Figs. 2D and 2E, Supplementary Fig. S2 and Table S4).

**Fig. 2.**
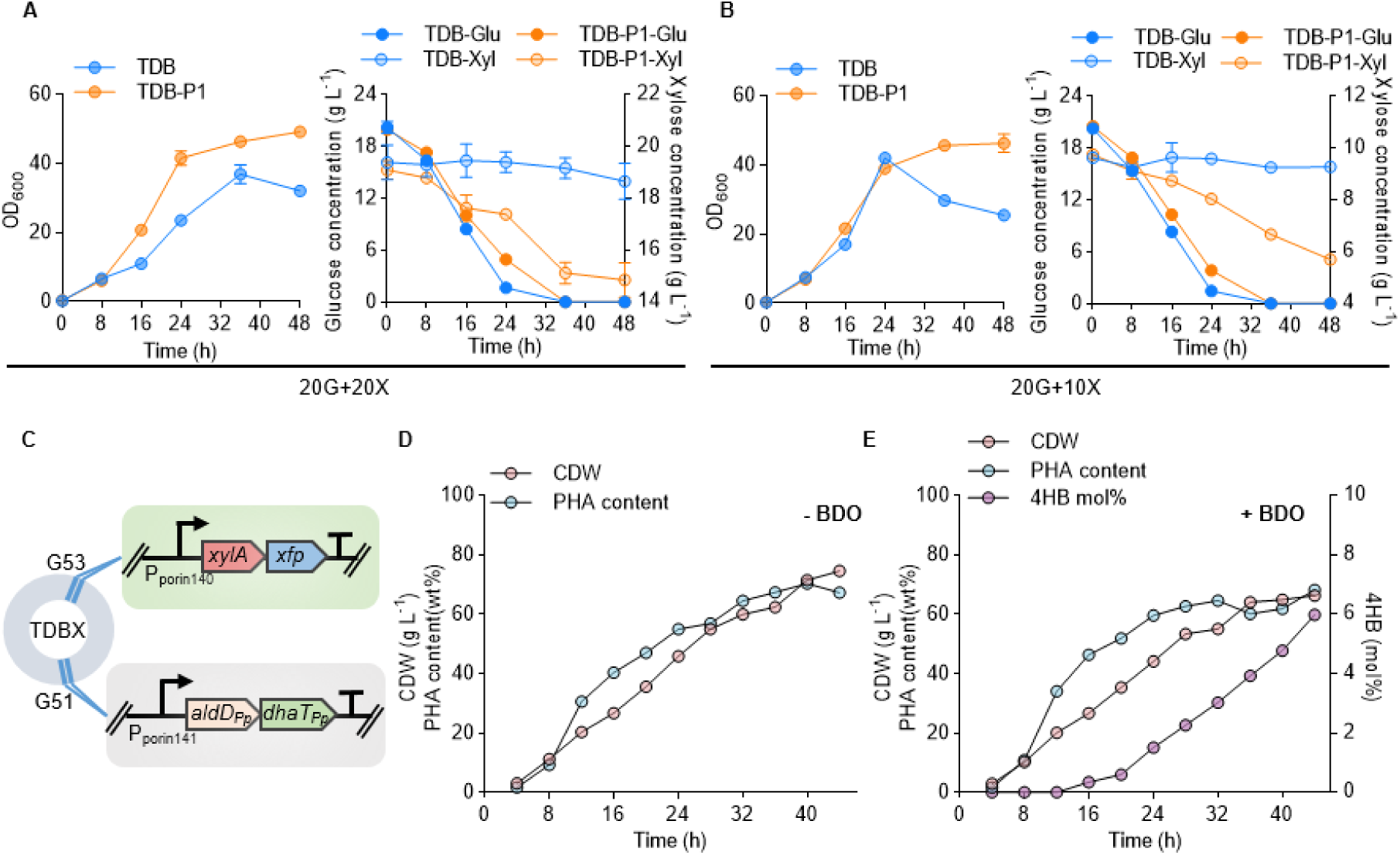
Synergetic co-utilization of glucose and xylose by engineered *Halomonas* TD. (**A**, **B**) Growth (OD_600_, left panel) and sugar-consumption (residual glucose and xylose, right panel) curves characterization by recombinant TDB carrying IPTG-induced (200 mg L^−1^) *xylA-xfp* module (TDB-P1) grown on mixed carbon sources (20G + 20X in part **A** and 20G + 10X in part **B**) compared to TDB strain under the same condition throughout the 48-h shake flask study. (**C**) Schematic design for constructing TDBX strain. The *xylA*-*xfp* gene cluster driven by P*_porin_*_140_ promoter was integrated on G53 loci based on TDB strain. (**D**, **E**) Fed-batch studies of cell dry weight (CDW) and PHA content (PHB in **D**, P34HB in **E**) by strain TDBX co-fed with glucose and xylose were conducted in a 7-L bioreactor. Specifically, 1,4-butanediol (BDO), the precursor for 4HB monomer synthesis, was added after 12-h cultivation for P34HB synthesis (**E**). Error bars represent standard deviations, n = 3, in part **A** and **B**; fed-batch studies in part **D** and **E** were carried out in single batch, n = 1.

In summary, the co-utilization of glucose and xylose in TDB strain has been achieved by engineering the phosphoketolase pathway to provide proven foundations and feasibility for converting renewable non-grain lignocellulosic hydrolysate into value-added biopolyesters by recombinant *Halomonas* chassis.

### 3.2. De novo synthesis of dipolymer P34HB from glucose and xylose

The P34HB synthesis from glucose and xylose has been succeeded by chromosomally engineered TDBX strain harboring fine-tuned phosphoketolase pathway, however, still depending on the supplementation of BDO used as a precursor for 4HB monomer accumulation, which has significant impact on the cell growth and P34HB material properties (e.g., tensile strength). Previous studies have demonstrated that the engineered *Halomonas* TD can accumulate P34HB from glucose only by redirecting the tricarboxylic acid (TCA) cycle flux (α-ketoglutarate and succinyl-CoA) towards 4HB synthesis^[49]^, which also can be employed to generate P34HB production by engineered TDBX using glucose and xylose only instead of BDO addition (Fig. 3A). Therefore, the previously fine-tuned de novo synthesis module of 4HB, namely *4hbd-sucD-ogdA* controlled by P_porin194_, was integrated on G49 locus of both TDBX and TDB strain after the deletion of *gabD*_1/2/3_ genes to form TDBGX and TDBG strain, respectively (Fig. 3B, Supplementary Fig. S3). Expectedly, over 6.5 g L^−1^ P(3HB-*co*- 5.6 mol% 4HB) was obtained by TDBGX grown on ‘20G+10X’ (Fig. 3C), which showed similar performance compared to the two TDBG recombinants harboring plasmid-carried *xylA*-*xfp* module controlled by P_MmP1_ (induced by 200 mg L^−1^ IPTG) and P_porin58_ (Supplementary Fig. S3), but significantly higher than TDBG strain producing only 4.7 g L^−1^ P34HB with 4.8 mol% 4HB, almost half decrease of 4HB mass accumulation (Fig. 3D), when grown on the same condition. This is mainly attributed to sufficient xylose utilization capability of TDBGX, consuming 7.2 g L^−1^ xylose, in contrast to TDBG unable cannot use xylose except to consuming the 20 g L^−1^ glucose.

**Fig. 3.**
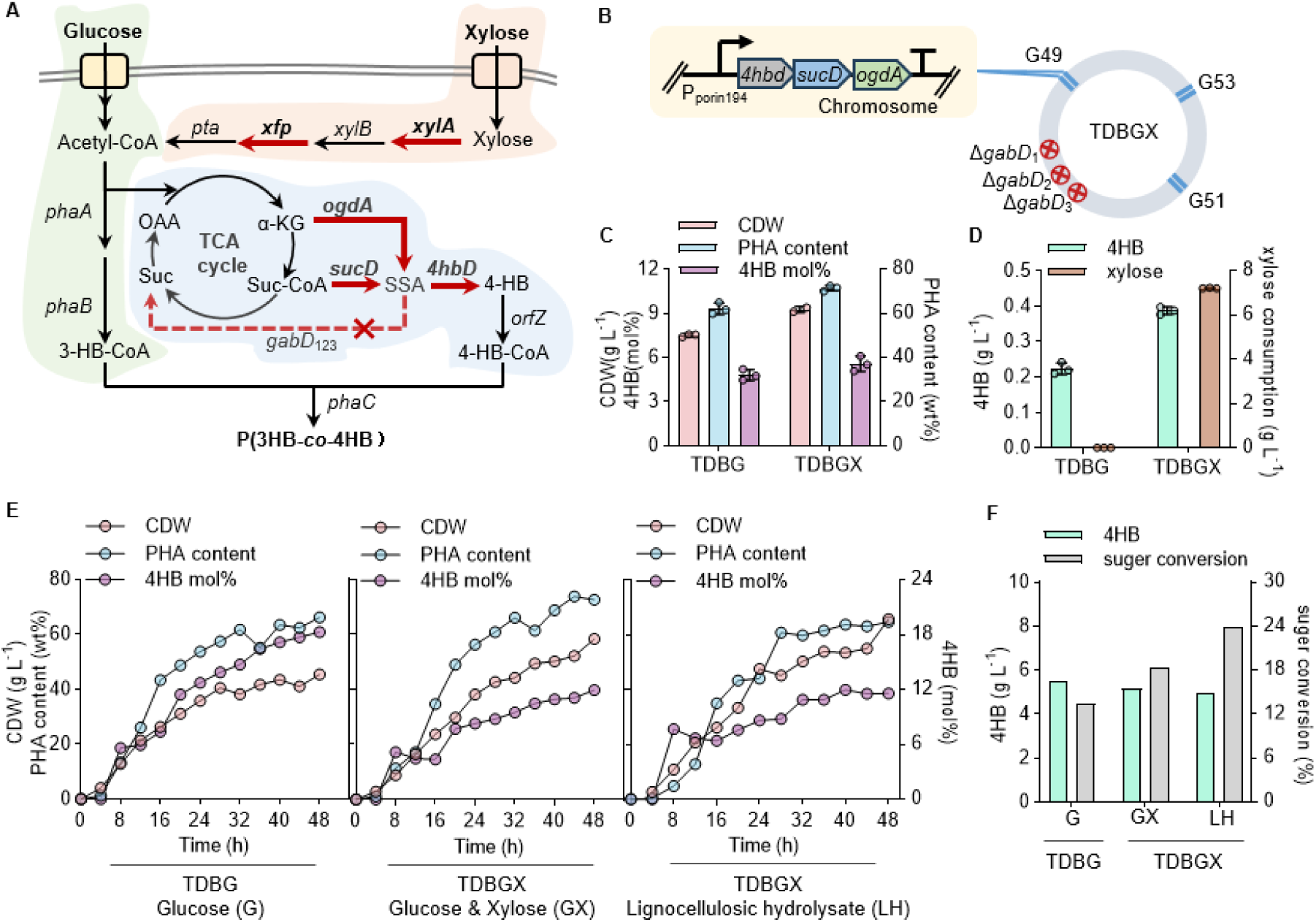
Engineering the de novo synthesis of P34HB from glucose and xylose. (**A**) Metabolic pathway design for de novo P34HB synthesis from glucose and xylose. Key enzymes and metabolites: *xylB* encoding xylulokinase; *pta* encoding phosphate acetyltransferase; *ogdA* encoding 2-oxoglutarate decarboxylase; *sucD* encoding succinate semialdehyde dehydrogenase; *4hbd* encoding 4-hydroxybutyrate dehydrogenase; *gabD*_1/2/3_ encoding three-copy succinate semialdehyde dehydrogenase; *phaA* encoding β-ketothiolase; *phaB* encoding NADPH-dependent acetoacetyl-CoA reductase; OAA, oxaloacetate; α-KG, α-ketoglutarate; Suc, succinate; Suc-CoA, succinyl-CoA; SSA, succinate semialdehyde. (**B**) Schema of constructing TDBGX strain for P34HB synthesis. The *4hbd*-*sucD*-*ogdA* cluster driven by P*_porin_*_194_ promoter was chromosomally integrated on G49 loci in TDBX after deleting the *gabD*_1/2/3_ genes to achieve reinforced 4HB synthesis flux. (**C**) Shake flask study of CDW, PHA content and 4HB molar ratio by TDBGX in contrast to TDBG, a derivate of TDB with defected *gabD*_1/2/3_ genes and containing *4hbd*-*sucD*-*ogdA* cluster on loci 53, after 48-h fermentation in 50 MM medium containing a mixed carbon source of ‘20G + 10X’. (**D**) The consumption of xylose and yield of 4HB monomer from part (**C**). (**E**) Fed-batch studies of P34HB production, including CDW, PHA content and 4HB molar ratio, were conducted in a 7-L bioreactor by recombinant TDBG using glucose only, and TDBGX using glucose and xylose (GX, middle panel) and lignocellulosic hydrolysate (LH, right panel), respectively. (**F**) Comparative analysis of sugar-to-PHA conversion rate and 4HB monomer yield after 48-h fermentation from part **E**. Error bars represent standard deviations, n = 3, in part **C** to **D**; fed-batch studies in part **E** were carried out in single batch, n = 1.

To assess the industrial scale-up feasibility, fed-batch fermentation for P34HB production by TDBGX utilizing mixed carbon source (glucose and xylose, GX group) and non-grain lignocellulosic hydrolysate (LH) from Juwei Yuanchuang Biotechnology Co., Ltd. (containing 408 g L^−1^ glucose and 174 g L^−1^ xylose analyzed by HPLC, LH group) was conducted in a 7 L bioreactor compared to TDBG grown in the same condition except using glucose as a sole carbon source (Fig. 3E, Supplementary Fig. S4). After 48 h cultivation, 42.6 g L^−1^ P(3HB-*co*-11.4 mol% 4HB) and 42.9 g L^−1^ P(3HB-*co*- 11.4 mol% 4HB) were obtained by TDBGX grown on mixed carbon source (GX) and LH, respectively, which were 40% higher than that by TDBG yielding only 30 g L^−1^ P(3HB-*co*- 18.2 mol% 4HB) (Fig. 3E, Supplementary Table S5). Additionally, although the 4HB mass accumulation level was of negligible variance across the three groups, higher sugar conversion rate could be achieved in both GX and LH group by TDGBX compared to TDGB grown on glucose only (Fig. 3F). Hence, the joint forces of introducing phosphoketolase pathway and de novo 4HB synthesis pathway in engineered TDB can lead to significant improvement on P34HB titer when co-utilizing glucose and xylose. Furthermore, the direct use of commercial non-grain LH by TDBGX exhibit no observed effect on P34HB accumulation, which indicates strong and robust metabolic capacity of xylose towards P34HB synthesis capable of efficient and cost-effect valorization of lignocellulosic resources by engineered *Halomonas* cells in the coming future.

### 3.3 De novo synthesis of terpolymer P34HBV from glucose and xylose

In addition to the phosphoketolase pathway-dependent xylose metabolism, which was successfully constructed in start host TDB for PHB and P34HB synthesis, another well-studied xylose metabolism mediated by Weimberg pathway is also important and worth studying. Since the identified Weimberg pathway from *Caulobacter crescentus* was reported able to convert xylose into BDO (C4)^[39]^ and α-ketoglutarate (C5)^[56]^, which can be potentially converted into 4HB-CoA and 3HV-CoA, respectively, to achieve diversified PHAs synthesis consisting of 4HB and/or 3HV monomers together with 3HB, including PHB, P34HB and PHBV copolymers, P34HBV terpolymer (Fig. 4A). Specifically, according to the previous reports, the 2-ketoacid decarboxylase (KDC) encoded by *kivD* from *Lactococcus lactis* can not only convert the xylose-derived intermediate 2,5-dioxopentanoate (DOP) into butanedial^[39]^, which was then converted into BDO by KDC (KviD) and alcohol dehydrogenase encoded by *yqhD*, but also convert α-ketoglutarate (α-KG) into 2-ketobutyrate (2-KB)^[57]^, which was then converted into propionyl-CoA for 3HV-CoA synthesis by wildtype *Halomonas* TD^[58]^. These previous findings indicate that the xylose-to-PHA diversity can be further expanded by designing and assembling different plug-and-play modules of xylose metabolism.

**Fig. 4.**
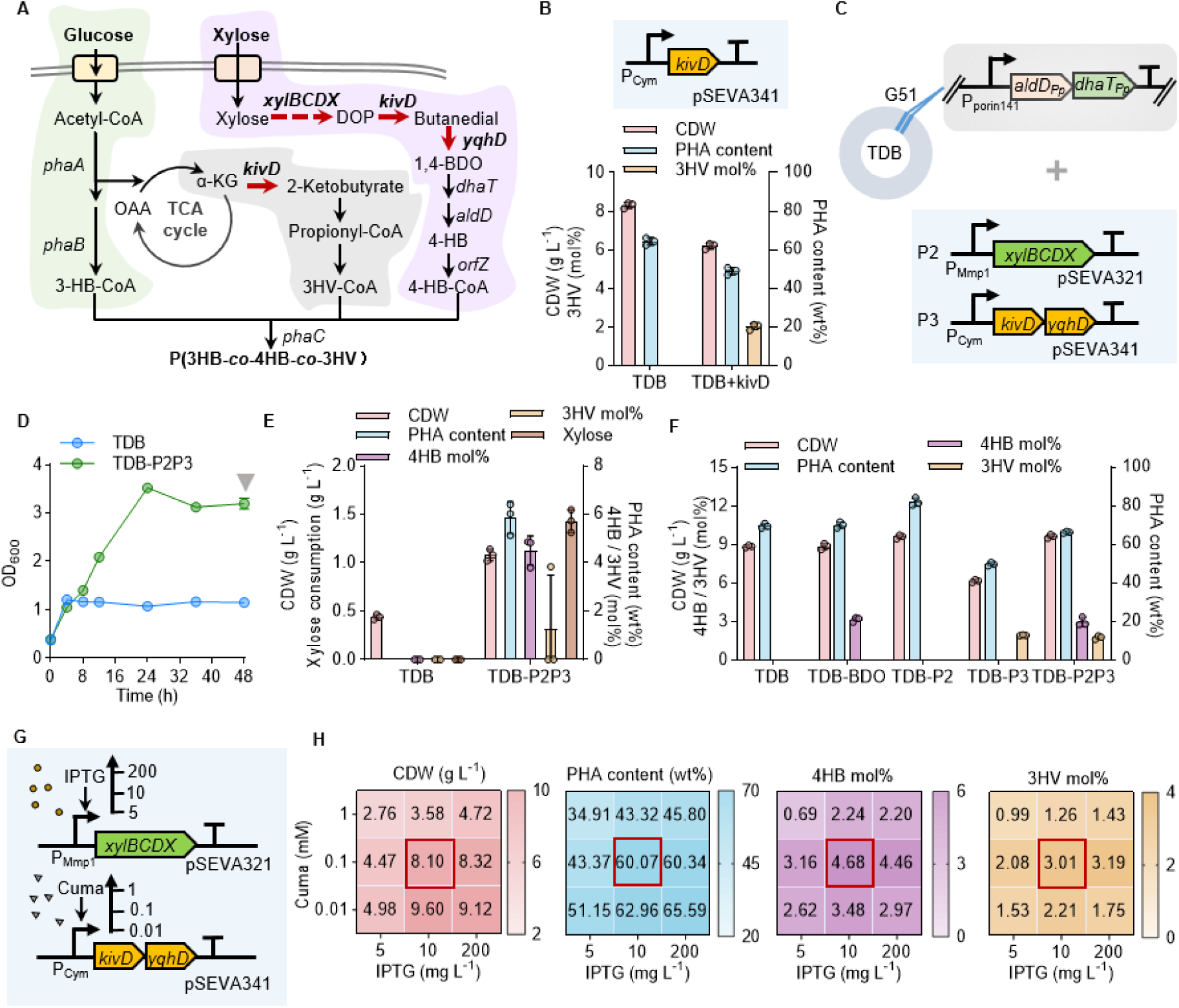
Engineering the de novo synthesis of terpolymer from glucose and xylose. (**A**) Metabolic pathway design for P(3HB-*co*-4HB-*co*-3HV) terpolymer, namely P34HBV, synthesis from glucose and xylose. Three enzymes encoded by *xylBCDX*, *kivD* and *yqhD* were employed to convert xylose into 4HB and 3HV monomers (arrows in red). Key enzymes and metabolites: *xylBCDX* gene cluster encoding xylose dehydrogenase, xylonolactonase, xylonate dehydratase and 2-keto-3-deoxy-xylonate dehydratase, respectively; *kivD* encoding keto-acid decarboxylase; *yqhD* encoding alcohol dehydrogenase; DOP, 2,5-dioxopentanoate. (**B**) Validation of 3-HV monomer (within PHBV) synthesis from glucose only by recombinant TDB harboring *kivD* expression module induced by cumic acid (0.1 mM). (**C**) Schema for terpolymer synthesis of P34HBV by recombinant TDB carrying two plasmids (TDB-P2P3): P2, containing *xylBCDX* cluster controlled by P_Mmp1_ induced by IPTG; P3, carrying *kivD-yqhD* cluster driven by P_Cym_ induced by cumic acid. (**D**) Cell growth assessment of TDB and TDB-P2P3 grown in 50 MM medium containing 10 g L^−1^ xylose in the presence of 200 mg L^−1^ IPTG and 0.01 mM cumic acid compared to TDB grown in the same condition without inducers. (**E**) Assessments of the CDW, PHA content, 4HB/3HV ratio (mol%), and xylose consumption by TDB and TDB-P2P3 grown on xylose only (sampled from part **D** with inverted triangle in gray). (**F**) Assessments of the CDW, PHA content, and monomer ratio (mol%) of 4HB and/or 3HV by TDB (PHB, P34HB) and its derivates harboring P2 and/or P3 grown on mixed carbon source containing 20 g L^−1^ glucose and 10 g L^−1^ xylose (20G + 10X). (**G**) Expression tuning design of *xylBCDX* and *kivD-yqhD* clusters using a dual-induction system (IPTG and cumic acid). (**H**) Comparative analysis of CDW, PHA content, and 4HB/3HV mol% by recombinant TDB-P2P3 grown on ‘20G + 10X’. All experiments were carried out with three parallels. Error bars represent standard deviations, n = 3.

To validate the α-KG-to-2-KB biocatalytic activity encoded by *kivD* for 3HV monomer synthesis, the expression module of *kivD* gene driven by a cumic acid (CA)-inducible system (P_Cym_) was firstly constructed on a plasmid-carried system and then conjugated into TDB strain (TDB-*kivD*). Compare to the start host TDB grown on 20 g L^−1^ glucose (20G) only harvesting PHB production, the TDB-*kivD* recombinant could produce PHA copolymer containing 98 mol% 3HB and 2 mol% 3HV under the same growth condition except for the supplementation of 0.1 mM CA (Fig. 4B). However, significant decreases of CDW and PHA content (CDW: 8.3 g L^−1^ *vs* 6.2 g L^−1^; PHA content: 64 wt% *vs* 48 wt%) were observed probably due to the overexpression of *kivD* gene or CA addition. Subsequently, the complete xylose-to-BDO biosynthesis pathway was constructed on two plasmids, namely P2 and P3 containing *xylBCDX* cluster controlled by IPTG-induced system (P_MmP1_, 200 mg L^−1^ IPTG) on pSEVA321 and *kivD-yqhD* cluster controlled by CA-induced system (P_Cym_, 0.1 mM CA) on pSEVA341, respectively (Fig. 4C). Shake flask study revealed that the terpolymer P34HBV, which was consisting of 94.2 mol% 3HB, 4.5 mol% 4HB and 1.3 mol% 3HV, could be successfully achieved by TDB-P2P3, TBD-derived recombinant containing P2 and P3 constructs, grown on 10 g L^−1^ xylose (10X). By contrast, poor growth and non-detectable PHA were observed by TDB grown on the same condition (Figs. 4D-4E). Moreover, different recombinants, including TDB and its derivates containing P2, P3 and P2+P3, respectively, grown on ‘20G+10X’ mixed carbon source were conducted in a 500-mL shake flask with an aim to obtain improved cell growth with enhanced PHA accumulation (Fig. 4F). Results showed that the introduction of P2 (Weimberg pathway) in TDB (TDB-P2) led to obvious improvement on CDW and PHA content compared to TDB group (CDW: 9.4 g L^−1^ *vs* 8.8 g L^−1^; PHA content: 82 wt% *vs* 69 wt%). However, the introduction of 2-ketoglutarate semialdehyde dehydrogenase (KSH) aiming to direct DOP flux into TCA cycle exhibited negative effects on both cell growth and PHA accumulation (Supplementary Fig. S6), which was not included in further study. Besides, similar to previous test (Fig. 4B), only P3HBV copolymer could be generated by recombinant TDB harboring P3 construct only (TDB-P3), and obvious decrease of CDW and PHA content were observed compared to TDB group (Fig. 4F). More importantly, the joint forces of P2 and P3 contained by TDB (TDB-P2P3) indeed boosted the terpolymer synthesis efficiently, yielding 6.3 g L^−1^ P(3HB-*co*- 3 mol% 4HB-*co*- 1.7 mol% 3HV) with similar levels of PHA titer, 4HB and 3HV molar ratio when compared to TDB (6.1 g L^−1^ PHB), TDB-BDO (3 mol% 4HB, supplementing 5 g L^−1^ BDO) and TDB-P3 (1.9 mol% 3HV) group, respectively (Fig. 4F, Supplementary Fig. S5).

To further elevate the 4HB and 3HV molar ratio within P34HBV terpolymer for improved material property, generally 5-to-10 mol% for non-3HB monomers (MedPHA Biotechnology Co., Ltd., https://www.medpha.cn/), expression tuning of *xylBCDX* and *kivD-yqhD* clusters was performed by TDB-P2P3 recombinant grown in different combinatory concentrations of IPTG (5, 10, and 200 mg L^−1^) and CA (0.01, 0.1, and 1 mM) (Fig. 4G). Although the highest CDW (9.6 g L^−1^) and PHA content (63 wt%) were achieved under the induction of 10 mg L[¹ IPTG and 0.01 mM CA (’10+0.01’), 10 mg L[¹ IPTG and 0.1 mM (‘10+0.1’) was identified to be the optimal induction combination, which could yield highest level of 4HB and 3HV molar ratio (∼ 7.7 mol% 4HB and 3HV in total) with only slightly decrease of CDW (8.1 g L^−1^) and PHA content (60 wt%) compared to ‘10+0.01’ group (Fig. 4H, Supplementary Table S6). Additionally, compared to the optimal group (’10+0.1’), even higher IPTG induction concentration (200 mg L^−1^) did not lead to any improvements on cell growth and terpolymer synthesis. In conclusion, the proposed bifunction of KivD and de novo synthesis of terpolymer P34HBV from glucose and xylose were successfully validated by engineered TDB harboring the Weimberg pathway-dependent xylose utilization (from xylose to DOP) cascaded with the shared pathway capable of simultaneous 3HV and 4HB monomer synthesis.

### 3.4 Enhanced P34HBV synthesis by developing self-induced xylose uptake system

As a co-utilized feedstock with glucose, the transmembrane transportation and intracellular metabolism generally determine the utilization efficiency of xylose. Herein, in addition to combining the phosphoketolase and Weimberg pathway for enhanced xylose metabolism, a xylose-induced system was designed and employed to control the expression of xylose transmembrane transporter with an aim to further boost the xylose uptake efficiency during the fed-batch fermentation once the residual xylose concentration exceeds a certain level (Fig. 5A). Firstly, a prototyping xylose-induced system was constructed consisting of a repressor protein XylR from *Bacillus subtilis*^[59]^ (sensing panel) and xylose-responsive promoter P_xylA_ using sfGFP as a reporter (control panel). The dose-response characterization was assessed by recombinant TDB harboring the xylose-induced system in the presence of 0, 1, 2, 3,5 and 10 g L^−1^ xylose. The highest induction level of normalized fluorescence (FI/OD_600_) reached up to 130 at 5 g L^−1^ xylose, however, obvious leakiness was observed with FI/OD_600_ value reaching up 25, which cannot be used for exquisite gene expression control directly (Fig. 5B). For a self-responsive induced system of xylose, low leakiness, high sensitivity (in response to lower control limit of feedstock), and wide-and-stable saturation induction range are of great significance to achieve robust transmembrane substrate-uptaking control during fermentation because of the inevitable fluctuation of residual sugar concentration (Fig. 5C).

**Fig. 5.**
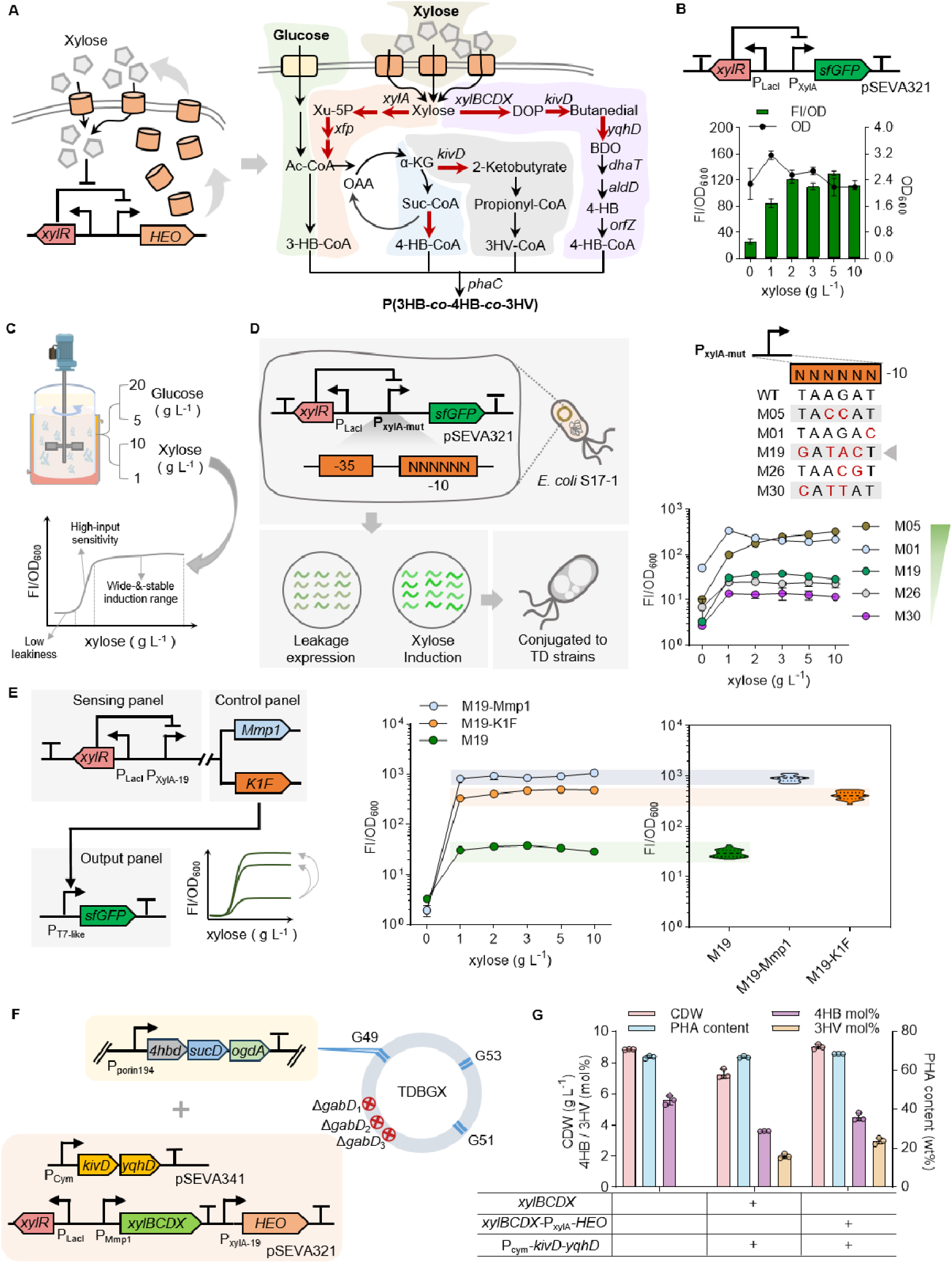
Enhancing the transmembrane transportation efficiency of xylose for improved terpolymer synthesis. (**A**) Schematic design for improved P(3HB-*co*-4HB-*co*-3HV) synthesis from glucose and xylose by enhancing the xylose transmembrane transportation efficiency. (**B**) Biosensor design and dose-response characterization of xylose-induced system using sfGFP as a reporter. (**C**) Schema for designing an xylose-control expression system suitable for practical uses during industrial fermentation process when using LH (glucose-to-xylose ratio ≈ 2:1) as a carbon source. The residual concentrations of glucose and xylose for PHA synthesis are generally from 5 g L^−1^ to 20 g L^−1^ and 1 g L^−1^ to 10 g L^−1^, which requires high-sensitive and stable expression output under wide range xylose concentration (1-10 g L^−1^), and low leakiness in the absence of xylose. (**D**) Library construction and screening workflow for desired xylose-induced system via random mutagenesis of P_xylA_ promoter (–10 region) (left panel). The best-performing mutant P_xylA-19_ (M19, inverted triangle) exhibiting tight leakiness and expected functions (medium output for transmembrane protein expression, high xylose sensitivity from 0-to-1 g L^−1^, wide and stable induction range) was obtained (right panel). (**E**) Characterization of the M19 xylose-control performance by cascading T7-like RNA polymerases, MmP1 and K1F that are similar to transmembrane protein like HEO requiring exquisite expression control, to validate the tight leakiness and wide-and-stable induction output under an over 10-fold amplified output. (**F**) Schematic design for enhanced terpolymer synthesis by recombinant TDBGX, which contains three chromosome-carried expression modules including *xylA-xfp* (G53), *aldD-dhaT* (G51) and *4hbd-sucD-ogdA* (G49), and two plasmid-carried modules including xylose transmembrane transporter (HEO) and xylose-to-4HB/3HV conversion pathway (*kivD-yqhD* and *xylBCDX*). (**G**) Comparative analysis of CDW, terpolymer [P(3HB-*co*-4HB-*co*-3HV)] content (wt%), 4HB and 3HV ratios (mol%) by TDBGX and its recombinants grown on glucose and xylose ‘20G + 10X’. FI/OD_600_, normalized fluorescent intensity (a.u.) by dividing OD_600_. Cell cultures were obtained and analyzed using a microplate reader (diluted to 0.2-0.8 of OD_600_). Error bars represent standard deviations, n = 3.

Therefore, random mutagenesis of the core region (–10 region) of P_xylA_ promoter was performed to obtain a mutated P_xylA_ library of varied induction strength using *E. coli*. S17-1 as a pre-testing host. The candidate mutants of tight leakiness were selected for further characterization in TDB to generate desired dose-response induction pattern (Figs. 5C and 5D, Supplementary Fig. S7A). Expectedly, two mutants, namely M05 (P_xyl-05_) and M19 (P_xyl-19_), exhibiting weaker basal fluorescence level (FI/OD_600_ = ∼10) and higher ON/OFF foldchange (> 10) were screened out (Fig. 5D, Supplementary Fig. S7B). Specifically, in contrast to the gradually rising induction level (FI/OD_600_) of M05, from 10 to 323, under increasing xylose dosage, the M19 mutant not only displays stable induction output with FI/OD_600_ value maintaining approximately at 30-to-38 when xylose concentration reaches no less than 1 g L^−1^, but also high sensitivity to low xylose concentration (∼1 g L^−1^). This mutant is an optimal candidate for xylose transmembrane up-taking control because the overexpression of most transmembrane transporters is toxic to engineered cells. Moreover, two amplifier systems were constructed based on M19 cascaded with different T7-like systems (MmP1 and K1F)^[44, 60]^ to evaluate the ON-/OFF-control robustness even under over 10-fold amplified output signal (Fig. 5E), which is also useful for genes requiring strong expression level like *phaCAB* cluster encoding PHB synthesis^[49]^. Notably, similar induction pattern could be observed in both 19-MmP1 and M19-K1F groups with lower leakiness (FI/OD_600_ < 2) and saturation output deviation (narrower color bar). These results indicate excellent control performance of M19 system with desired function.

Therefore, a self-induced xylose uptake module, which contains a previously reported xylose transporter encoded by gene *HEO* from *Halomonas elongata*^[51, 61]^ driven by P_xyl-19_, was constructed on P2 plasmid, namely P4. The shake flask results showed obvious improvement on CDW (7.3 g L^−1^ *vs* 9.0 g L^−1^), 4HB (3.6 mol% *vs* 4.5 mol%) and 3HV (2 mol% *vs* 3 mol%) molar ratio within P34HBV by recombinant TDBGX harboring P3 and P4 plasmids (TDBGX-P3P4) compared to control group by TDBGX-P2P3 (Fig. 5F). Besides, in contrast to TDGBX group producing P34HB, the joint carbon flux of 4HB and 3HV (∼7.5 mol%) was obvious higher than that of 4HB in P34HB (∼5.6 mol%). These results demonstrated proven success on improving xylose utilization efficiency by introducing self-induced uptake system following the rationale of begin-with-the-end-in-mind design.

## 4. Conclusions

This study successfully constructed different recombinant *Halomonas* strains able to utilize glucose and xylose, or lignocellulose hydrolysate, as feedstock efficiently for various PHAs synthesis, including PHB (Fig. 2D), P(3HB-*co*-4HB) (Fig. 3E) and P(3HB-*co*-4HB-*co*-3HV) (Figs. 4-5), by introducing the phosphoketolase and/or Weimberg pathways generally encoding the xylose metabolism. More importantly, the fed-batch study of effective P34HB production by TDBGX recombinant grown on commercial lignocellulosic hydrolysate (LH, purchased from Juwei Yuanchuang Biotechnology Co., Ltd) demonstrated strong practical value and industrialized feasibility for lignocellulose upcycling (Fig. 3E). Certainly, more attempts focusing on process optimization of fed-batch fermentation can be employed to achieve further improvement on cell growth and PHA accumulation.

As a rising-star chassis allowing unsterile and open fermentation, although *Halomonas* TD has already been engineered to use xylose as a carbon source in previous studies^[51, 53]^, the diversity of PHA production was limited but further expanded in this study. For example, engineering extended Weimberg pathway that converts xylose into BDO, a precursor intermediate for 4HB monomer accumulation, was ingeniously designed to achieve copolymer synthesis consisting of 3HB and 4HB (Fig. 4A). Meanwhile, the proposed bifunction of keto-acid decarboxylase encoded by *kivD*, which can convert butanedial and α-ketoglutarate into BDO and 2-ketobutyrate, respectively, was validated and used to obtain P34HBV production, a terpolymer of outstanding material property^[22]^. Notably, these results also provide an alternative pathway design for 3HV monomer synthesis derived from TCA-cycle compound (α-ketoglutarate). Moreover, a well-designed xylose biosensing system was developed and optimized based on *Halomonas* TD for tailoring robust and self-induced xylose uptake during the fermentation process, which led to obvious improvement on terpolymer P(3HB-*co*-4HB-*co*-3HV) synthesis incorporating the joint forces of phosphoketolase and Weimberg pathway-based xylose utilization, including over 23%, 25% and 50% increase of CDW, 4HB and 3HV molar ratios (Fig. 5G), respectively, by engineered TDBGX grown on glucose and xylose compared to the control groups.

In addition to the achievements on xylose utilization and PHA synthesis, more efforts are still required for the upcoming study focusing on lignocellulosic biomass utilization based on recombinant *Halomonas*: I) employing AI-assisted tools and methods to mine and design enzymes related to xylose metabolism for more efficient xylose utilization (e.g. enhancing the cell growth when using xylose as feedstock from Fig. 2D) and PHA synthesis (e.g. achieving independent or ratio-controllable synthesis of 3HV and 4HB by engineered KivD from Fig, 4A); II) developing neutral high-expression chromosomal loci for multi-module pathway integration to construct strain allowing antibiotic-free and robust fermentation process of industrial purpose (e.g. obtaining chromosomal expression of self-induced xylose uptake system and Weimberg pathway from Fig. 5F); III) constructing whole-genome model-based carbon metabolic network for directing reinforced flux towards PHA synthesis (e.g. improving the substrate-to-PHA conversion rate from Fig. 3F); etc. Therefore, the implementation of these strategies would further strengthen the feasibility of industrialized PHAs production by recombinant *Halomonas* using xylose or substrate containing xylose.

In conclusion, this study demonstrates strong potential and significance of engineering *Halomonas* for both lignocellulosic resource valorization and non-grain-dependent PHA manufacturing.

## Author contributions

**Chang-Le Shen:** Writing-original draft, Visualization, Resources, Methodology, Investigation, Formal analysis, Data acquisition and curation**. Bao-Wen Liu:** Methodology, Resources, Formal analysis, Data acquisition and curation**. Peng Liu:** Resources, Formal analysis, Data acquisition and curation**. Yi-Hao Deng:** Resources, Formal analysis. **Ling-Ling Zhao:** Resources. **Liu-Song Yu**: Resources. **Wei Situ**: Resources. **Jin-Yan Lv**: Resources. **Hong-Wei Shen**: Resources. **Yi-Na Lin:** Resources, Methodology, Formal analysis, Data curation, Funding acquisition**. Yan-Chun Xiao:** Resources, Methodology, Formal analysis, Data curation. **Hai-Tao Yue:** Resources, Methodology, Formal analysis, Data curation**. Jian-Wen Ye:** Proposed the idea, designed and supervised the research, Writing-review & editing, Resources, Methodology, Validation, Formal analysis, Conceptualization, Supervision, Funding acquisition.

## Declaration of competing interest

The authors declare that they have no known competing commercial interests in this paper.

## Data availability

The data supporting this article have been included as part of the Supplementary Information. Supplementary information is available.

## Acknowledgments

We are thankful to Professor Guo-Qiang Chen (CGQ, Tsinghua University) for kindly donating the *Halomonas* TD01 strain. This research was financially supported by grants from National Natural Science Foundation of China (Grant No. 32322003 to YJW, No. 32501308 to LYN), the Fundamental Research Funds for the Central Universities (Grant No. 2025ZYGXZR096 to YJW) and Guangdong Basic and Applied Basic Research Foundation (No. 2026A1515011148 to LYN). We acknowledge the related fundings supported by the joint laboratory program of MedPHA Biotechnol. Co., Ltd and SCUT.

## Notes

### Competing Interest Statement

The authors have declared no competing interest.

