## Supporting Information for "Engineering xylose metabolism for diverse polyhydroxyalkanoates synthesis in *Halomonas* TD"

Supplementary Tables

**Table S1 Strains and genes used in this study.**

| **Strains** | **Descriptions** | **References** |
| --- | --- | --- |
| *E. coli* S17-1 | A vector donor used for conjugation harboring the *tra* genes from plasmid RP4 in the genome | Simon et al., 1983^[1]^ |
| TD01 | Wild-type *Halomonas* TD01 strain isolated from Aydingkol Lake in Xinjiang Province, China | Tan et al., 2011^[2]^ |
| TDH4 | TD01 derivate with defected *phaP*_1_ gene associated with PHA granule size control and integrated with *orfZ* gene encoding 4HB-CoA transferase | Shen et al., 2019^[3]^ |
| TDB | TDH4 derivate integrated with ‘P_porin141_-*aldD-dhaT*’ expression module on loci G51 | This study |
| TDBX | TDB derivate integrated with ‘P_porin140_-*xylA-xfp*’ expression module on loci G53 | This study |
| TDBG | TDB derivate integrated with ‘P_porin194_-*4hbd-sucD-ogdA*’ expression module on loci G49 together with defected *gabD*_1_, *gabD*_2_ and *gabD*_3_ genes | This study |
| TDBGX | TDBG derivate integrated with ‘P_porin140_-*xylA-xfp*’ expression module on loci G53 | This study |

**Table S2 Plasmids used in this study.**

| **Plasmid** | **Descriptions** | **References** |
| --- | --- | --- |
| pSEVA321 | Expression vector containing *oriT* for the expression of target genes in TD01, RK2 replication origin, Cm^R^. | Silva-Rocha et al., 2013^[4]^ |
| pSEVA341 | Expression vector containing *oriT* for the expression of target genes in TD01, pRO1600 replication origin, Km^R^ and Sp^R^. | Silva-Rocha et al., 2013^[4]^ |
| P1 | pSEVA321 derivative containing *xylA-xfp* cluster controlled by P_Mmp1_, Cm^R^. | This study |
| pP_porin58_-*xylA-xfp* | pSEVA321 derivative containing *xylA-xfp* cluster controlled by P_porin58_, Cm^R^. | This study |
| pP_Cym_-*kivD* | pSEVA341 derivative containing *kivD* module controlled by P_Cym_, Km^R^ and Sp^R^. | This study |
| P2 | pSEVA321 derivative containing *xylBCDX* cluster controlled by P_Mmp1_, Cm^R^. | This study |
| P3 | pSEVA341 derivative containing *kivD-yqhD* cluster controlled by P_Cym_, Km^R^ and Sp^R^. | This study |
| pP_Cym_-*KSH* | pSEVA341 derivative, *ksh* controlled by P_Cym_, Km^R^ and Sp^R^. | This study |
| pP_LacI_-*xylR-*P_xylA_-*sf*GFP | pSEVA321 derivative containing *xylR* controlled by P_LacI_ and *sf*GFP controlled by P_xylA_, Cm^R^. | This study |
| pP_LacI_-*xylR-*P_xylA(mutant)_-*sf*GFP | pP_LacI_-*xylR-*P_xylA_-*sf*GFP derivates based on the random mutagenesis of P_xylA_ (-10 region), Cm^R^. | This study |
| P4 | pSEVA321 derivative, *xylR* controlled by P_LacI_, *xylBCDX* controlled by P_Mmp1_, *HEO* controlled by P_xylA-19_, Cm^R^, pP_LacI_-*xylR-*P_Mmp1_*-xylBCDX-*P_xylA-19_-*HEO.* | This study |
| pP_LacI_-*xylR* P_xylA-19_-*Mmp1-*P_Mmp1_*-sf*GFP | pSEVA321 derivative, *xylR* controlled by P_LacI_, *Mmp1* controlled by P_xylA-19_, *sf*GFP controlled by P_Mmp1_, Cm^R^. | This study |
| pP_LacI_-*xylR* P_xylA-19_-*K1F-*P_K1F_*-sf*GFP | pSEVA321 derivative, *xylR* controlled by P_LacI_, *K1F* controlled by P_xylA-19_, *sf*GFP controlled by P_K1F_, Cm^R^. | This study |
| pP_Porin43_-*kivD-yqhD* | pSEVA341 derivative, *kivD-yqhD* controlled by P_Porin43_, Km^R^ and Sp^R^. | This study |
| pP_Porin185_-*kivD-yqhD* | pSEVA341 derivative, *kivD-yqhD* controlled by P_Porin185_, Km^R^ and Sp^R^. | This study |
| pP_Porin226_-*kivD-yqhD* | pSEVA341 derivative, *kivD-yqhD* controlled by P_Porin226_, Km^R^ and Sp^R^. | This study |
| pP_Porin12_-*kivD-yqhD* | pSEVA341 derivative, *kivD-yqhD* controlled by P_Porin12_, Km^R^ and Sp^R^. | This study |

**Table S3 Genes used in this study.**

| **Genes** | **Descriptions** | **References** |
| --- | --- | --- |
| *aldD* | Encoding aldehyde dehydrogenase from *Pseudomonas putida* KT2440, GenBank No.: AAN66172.1 | Lab stocks |
| *dhaT* | Encoding 1,3-Propanediol dehydrogenase from *Pseudomonas. putida* KT2440, GenBank No.: AAN68411.1 | Lab stocks |
| *xylA* | Encoding xylose isomerase from *Escherichia coli* MG1655 | Tan et al., 2022^[5]^ |
| *xfp* | Encoding phosphoketolase from *Lactobacillus rhamnosus* CGMCC1.120 | Tan et al., 2022^[5]^ |
| *4hbd* | Encoding 4-hydroxybutyrate dehydrogenase from *C. kluyveri* | Li et al., 2010^[6]^ |
| *sucD* | Encoding succinate semialdehyde dehydrogenase from *C. kluyveri Caulobacter* | Li et al., 2010^[6]^ |
| *ogdA* | Encoding 2-oxoglutarate decarboxylase from cyanobacterial, *Synechococcus sp.* PCC 7002 | Zhang et al., 2015^[7]^ |
| *gabD_x_* | Endogenous succinate semialdehyde dehydrogenase of *Halomonas.* TD01 | Ye et al., 2018^[8]^ |
| *xylB* | Encoding D-xylose dehydrogenase from *Caulobacter crescentus* NA1000 | Tai et al., 2016^[9]^ |
| *xylC* | Encoding D-xylonolactonase from *C. crescentus* NA1000 | Tai et al., 2016^[9]^ |
| *xylD* | Encoding D-xylonate dehydratase from *Caulobacter crescentus* NA1000 | Tai et al., 2016^[9]^ |
| *xylX* | Encoding 2-keto-3-deoxy-D-xylonate dehydratase from *C. crescentus* NA1000 | Tai et al., 2016^[9]^ |
| *kivD* | Encoding 2-ketoacid decarboxylase from *Lactococcus lactis* IFPL730 | Tai et al., 2016^[9]^ |
| *yqhD* | Encoding alcohol dehydrogenase from *E. coli* MG1655 | Tai et al., 2016^[9]^ |
| *ksh* | Endogenous aldehyde dehydrogenase of *Halomonas* TD01 | This study |
| *HEO* | Encoding xylose transporter from *Halomonas elongata* OUT30018 | Tan et al., 2022^[5]^ |
| *xylR* | Encoding xylose regulators from *Bacillus subtilis* | Wu et al., 2023^[10]^ |
| *mmp1* | Encoding T7-like RNA polymerase, MmP1 | Zhao et al., 2017^[11]^ |
| *k1F* | Encoding T7-like RNA polymerase, K1F | Zhao et al., 2017^[11]^ |

**Table S4 CDW, PHA content, 4HB mol% and sugar consumption by recombinant TDBX after 44-h fermentation conducted in a 7 L bioreactor.**

| **Strain** | **Sourced figure** | **CDW (g L^-1^)** | **PHA (wt%)** | **4HB (mol%)** | **Sugar consumption (g)** | |
| --- | --- | --- | --- | --- | --- | --- |
|  |  |  |  |  | **Glucose** | **Xylose** |
| TDBX (BDO-) | Fig. 2F | 74.5 | 67.2 |  | 826 | 169.2 |
| TDBX (BDO+) | Fig. 2G | 66.3 | 68.2 | 6 | 844 | 193.5 |

**Table S5 Fed-batch study of CDW, PHA content, 4HB mol%, and sugar consumption by recombinant TDBG and TDBGX, respectively, after 48-fermentation conducted in a 7-L bioreactor (from Fig. 3E).**

| **Strain** | **CDW (g L^-1^)** | **PHA (wt%)** | **4HB (mol%)** | **Sugar consumption(g)** | |
| --- | --- | --- | --- | --- | --- |
|  |  |  |  | **Glucose** | **Xylose** |
| TDBG | 45.4 | 66.2 | 18.2 | 1123 | / |
| TDBGX | 58.8 | 73.0 | 12.0 | 962 | 212 |
| TDBGX (LH) | 65.8 | 64.7 | 11.4 | 641.4 | 251 |

**Table S6 Shake flask study of CDW, PHA content, monomer ratio (mol%, 4HB and 3HV) based on the simultaneous tuning of *xylBCDX* and *kivD-yqhD* cluster from Figs. 4H.**

|  | **Cuma (mM)** | **IPTG (mg L^-1^)** | | |
| --- | --- | --- | --- | --- |
|  |  | **5** | **10** | **200** |
| **CDW (g L^-1^)** | **1** | 2.76±0.20 | 3.58±0.24 | 4.72±0.12 |
|  | **0.1** | 4.47±0.60 | 8.10±0.23 | 8.32±0.34 |
|  | **0.01** | 4.98±0.02 | 9.60±1.01 | 9.12±0.88 |
| **PHA content (wt%)** | **1** | 34.91±6.77 | 43.32±1.57 | 45.80±2.84 |
|  | **0.1** | 43.37±0.99 | 60.07±6.73 | 60.34±8.07 |
|  | **0.01** | 51.15±2.65 | 62.96±5.41 | 65.59±4.84 |
| **4HB (mol%)** | **1** | 0.69±0.18 | 2.24±0.16 | 2.20±0.38 |
|  | **0.1** | 3.16±0.01 | 4.68±0.55 | 4.46±0.15 |
|  | **0.01** | 2.62±0.27 | 3.48±0.61 | 2.97±0.34 |
| **3HV (mol%)** | **1** | 0.99±0.39 | 1.26±0.10 | 1.43±0.17 |
|  | **0.1** | 2.08±0.40 | 3.01±0.10 | 3.19±0.22 |
|  | **0.01** | 1.53±0.07 | 2.21±0.11 | 1.75±0.13 |

**Supplementary figures**

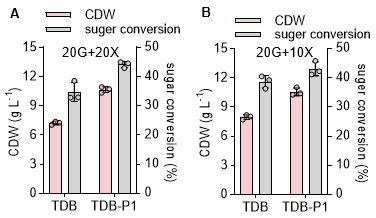

**Fig. S1 Comparative analysis of CDW and sugar conversion rate by recombinant *Halomonas* grown on glucose and xylose.**

(**A** and **B**) CDW and total sugar-to-CDW conversion rate by TDB strain and its recombinant TDB-P1 harboring plasmid-carried *xylA-xfp* grown on 20 g L^-1^ glucose + 20 g L^-1^ xylose (**A**, from Fig. 2A) and 20 g L^-1^ glucose + 10 g L^-1^ xylose (**B**, from Fig. 2B), respectively. Error bars represent standard deviations, n = 3.

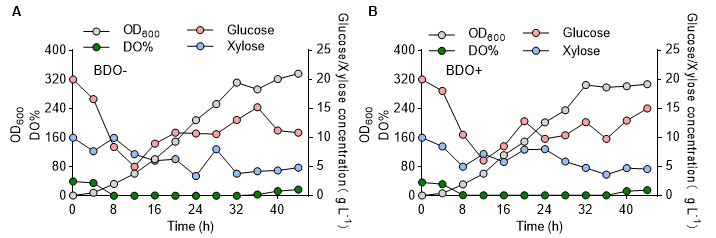

**Fig. S2 Time-course profiling of fed-batch studies by TDBX.**

Time-course profiling of OD_600_, dissolved oxygen (DO%) and residual concentrations of glucose and xylose during fed-batch studies by strain TDBX grown in a 7 L bioreactor without (**A**) and with (**B**) BDO addition (Figs. 2D and 2E), which is used as a precursor for 4HB monomer synthesis.

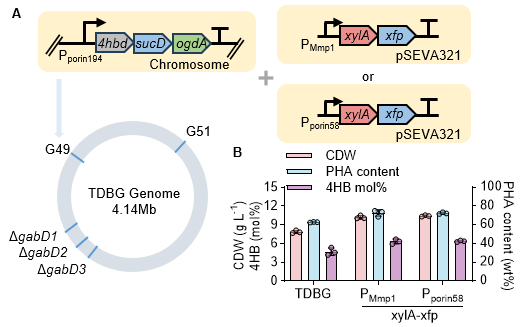

**Fig. S3 Engineering de novo synthesis of P34HB by recombinant TDBG.**

(**A**) Schematic design for P34HB synthesis by recombinant TDBG harboring *xylA-xfp* expression module controlled by P_MmP1_ or P_porin58_. The *4hbd-sucD-ogdA* cluster driven by P_porin194_ was chromosomally integrated on G49 loci in TDB after the deletion of gabD_1/2/3_ genes, forming TDBG. (**B**) Shake flask study of CDW, PHA content and 4HB mol% by TDBG and its derivatives after 48-h cultivation in 50 MM medium containing 20 g L^-1^ glucose + 10 g L^-1^ xylose (20G + 10X). For P_MmP1_ group, 200 mg L⁻¹ IPTG was initially added to the medium to induce the expression of *xylA-xfp*. Error bars represent standard deviations, n = 3.

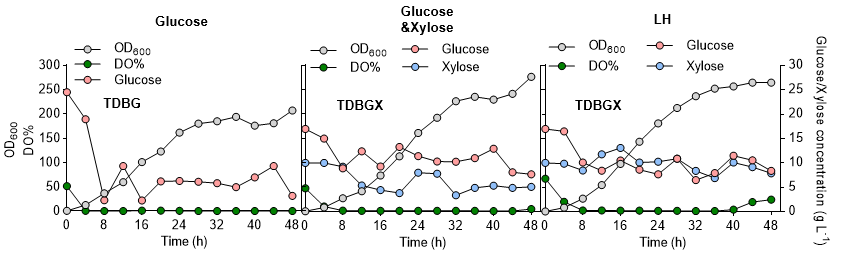

**Fig. S4 Time-course profiling of fed-batch studies for P34HB production by different recombinants.**

Time-course profiling, including OD_600_, DO% and residual concentrations of glucose and xylose, during fed-batch studies (from Fig. 3E) by TDBG grown on glucose only (left panel), and TDBGX grown on glucose and xylose (middle panel) and lignocellulosic hydrolysate from Juwei Yuanchuang Biotechnology Co., Ltd (LH group in right panel). Single fed-batch study, n = 1.

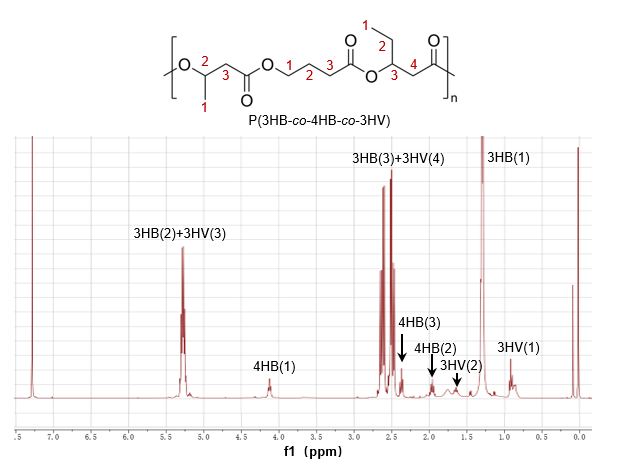

**Fig. S5 ^1^H NMR analysis of the obtained P(3HB-*co*-4HB-*co*-3HV) terpolymer.**

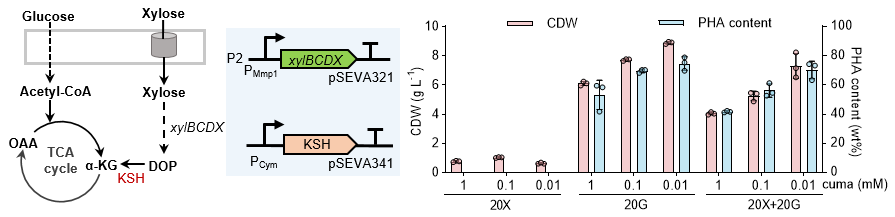

**Fig. S6 Testing the Weimberg pathway for xylose utilization in *Halomonas* TDB.**

The 2-ketoglutarate semialdehyde dehydrogenase (KSH) encoded by *ksh* can convert xylose-derived intermediate metabolite DOP into α-KG involved in TCA cycle. The construct containing *ksh* controlled by P*_Cym_* (induced by cumic acid) was conjugated into TDB strain together with P2 plasmid (10 mg L^-1^ IPTG) for shake flask fermentation using 50 MM media supplemented with 20 g L^-1^ xylose (20X), 20 g L^-1^ glucose (20G), and 20 g L^-1^ glucose + 20 g L^-1^ xylose (20G + 20X), respectively. Specifically, 0.01, 0.1, and 1 mM cumic acid were initially added to the medium, respectively, to achieve different induction level of KSH. CDW and PHA content of each testing group were measured after 48-h cultivation. Error bars represent standard deviations, n = 3.

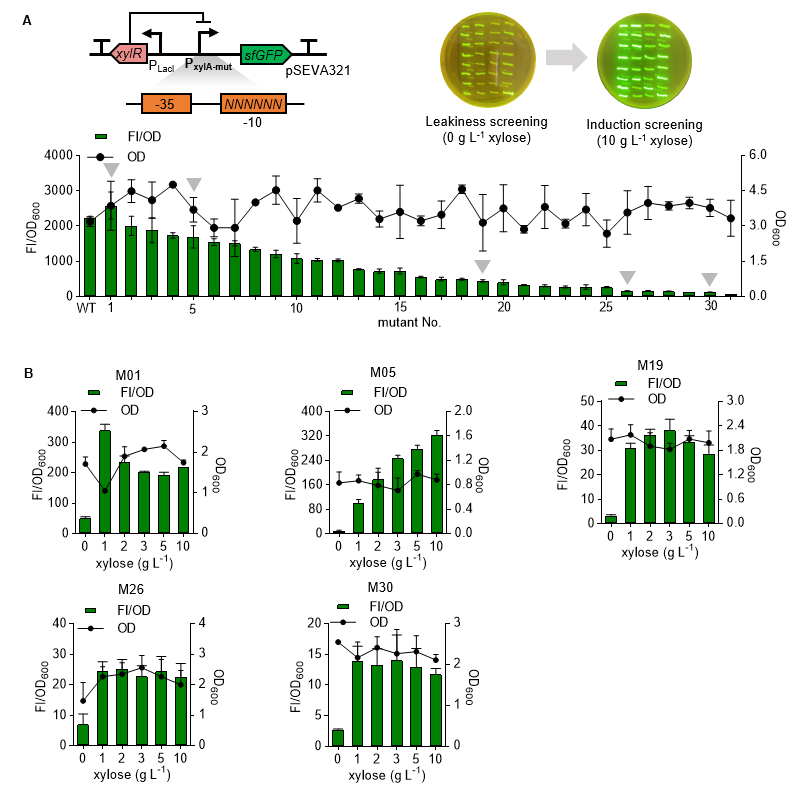

**Fig. S7 Construction and optimization of xylose-inducible system in *Halomonas*.**

(**A**) Characterization of expression levels by different P_xylA_ mutants in *E. coli* S17-1. Comparative analysis of different fluorescent cells (*E. coli* S17-1) harboring different P_xylA_ mutants grown on a LB agar plate supplemented with 10 g L^-1^ against the ones without xylose (0 g L^-1^). Fluorescence of different recombinant cells grown in LB medium containing 10 g L^-1^ xylose was measured by a microplate reader. (**B**) Dose-response characterization of different P_xylA_ mutants (M01, M05, M19, M26, M30) in *Halomonas* TDB. Fluorescence of different recombinant cells grown in 50MM medium containing 0, 1, 2, 3, 5 and 10 g L^-1^ xylose, respectively, was measured by a microplate reader. FI, Fluorescent intensity in arbitrary unit (a.u.). FI/OD_600_, normalized fluorescence by dividing OD_600_. Cell cultures were obtained and diluted to 0.2-0.8 of OD_600_ for FI/OD_600_ analysis using a microplate reader. Error bars represent standard deviations, n = 3.
